# HPC-REDItools: a Novel HPC-aware Tool for Improved Large Scale RNA-editing Analysis

**DOI:** 10.1101/2020.04.30.069732

**Authors:** Tiziano Flati, Silvia Gioiosa, Nicola Spallanzani, Ilario Tagliaferri, Maria Angela Diroma, Graziano Pesole, Giovanni Chillemi, Ernesto Picardi, Tiziana Castrignanò

## Abstract

**Background:** RNA editing is a widespread co-/post-transcriptional mechanism that alters primary RNA sequences through the modification of specific nucleotides and it can increase both the transcriptome and proteome diversity. The automatic detection of RNA-editing from RNA-seq data is computational intensive and limited to small data sets, thus preventing a reliable genome-wide characterisation of such process.

**Results:** In this work we introduce HPC-REDItools, an upgraded tool for accurate RNA-editing events discovery from large dataset repositories. Availability: https://github.com/BioinfoUNIBA/REDItools2.

**Conclusions:** HPC-REDItools is dramatically faster than the previous version, REDItools, enabling big-data analysis by means of a MPI-based implementation and scaling almost linearly with the number of available cores.

## Background

Advances in next generation sequencing (NGS) technologies have led to the production of an unprecedented amount of omic data (including genomes, transcriptomes, epigenomes from cells, tissues and organisms) changing science and medicine in ways never seen before and entering the “big data” era. The scale and efficiency of NGS poses the relevant challenges of sharing, archiving, integrating and analyzing these vast collections of omic data. Although tools and algorithms to handle and analyse large NGS datasets are now appearing, widespread software for common NGS analyses are yet multi-threaded or serial and not ready for the “big data” era revolution. As a consequence, the investigation of specific and important biological phenomena in large NGS datasets from international projects - such as The Cancer Genome Atlas (TCGA) [1]^[1]^ or The Genotype-Tissue Expression project (GTEx) [2]^[2]^ - is somehow precluded and, thus, a thorough redesign of algorithms and efficient implementations on High Performance Computing (HPC) infrastructures are mandatory [3]. Hereafter, we introduce and describe HPC-REDItools, a HPC tool for accurate detection of RNA-editing events from large data repositories.

RNA editing is a widespread post-transcriptional mechanism that alters primary RNA sequences through specific base modifications and increases the transcriptome complexity of eukaryotic organisms. In humans, RNA editing affects nuclear and cytoplasmic transcripts mainly by the deamination of adenosine (A) to inosine (I) through the ADAR family of enzymes [4] that acts on double RNA strands. Since I is commonly interpreted as guanosine by translation and splicing machineries (other than sequencing enzymes), A-to-I modifications can alter codon identity or base-pairing interactions within higher-order RNA structures [5]. As a result, A-to-I RNA editing can increase proteome diversity or regulate gene expression at the RNA level [5]. Moreover, editing within pre-mRNAs can generate or destroy splice sites, modulate alternative splicing and influence the dynamics of constitutive splice sites. A-to-I RNA editing is prominent in non-coding regions containing repetitive elements (mainly SINEs belonging to the Alu family) and rare in protein coding portions of genes [6, 7].

RNA editing has relevant and serious biological and physiological implications. Indeed, its deregulation has been linked to several nervous and neurodegenerative diseases such as epilepsy, schizophrenia, major depression, Alzheimer and amyotrophic lateral sclerosis [8]. In addition, the functional importance of this mechanism was established by showing that mice lacking ADARs die in utero or soon after weaning. Recently, editing alterations have also been associated with a variety of human cancers [9, 10].

Despite its importance in modulating gene expression and maintaining a correct cellular homeostasis, the A-to-I landscape in human is still incomplete and main biological roles are yet elusive. Indeed, the de novo detection of RNA editing in humans has been performed in a limited number of samples, tissues and experimental conditions.

Thanks to international consortia, thousands of transcriptome experiments (RNA-seq) have been performed and publically released through specialised web archives such as dbGAP (the database of Genotypes and Phenotypes [11]) or SRA (the Sequence Read Archive [12]). The Genotype-Tissue Expression (GTEx) consortium, for instance, provides the largest collection of RNA-seq experiments from 55 human healthy body sites of more than 900 individuals. RNA-seq collections like GTEx represent precious resources to investigate RNA editing in a multiplicity of human tissues.

In order to detect RNA editing sites in RNA-seq data, we developed the REDItools package [13, 14], a bioinformatics resource implemented in the portable Python programming language. Although one of the most accurate software for this purpose [15] and memory efficient (from 2 to 4 GB are generally sufficient), it is computationally intensive and time-consuming. In order to investigate the RNA editing landscape in very large cohort of RNA-seq datasets, we re-designed the main algorithm, optimizing its implementation for HPC infrastructures. The novel algorithm is on average 8-10 times faster than the previous version on a single core, while the HPC implementation scales almost linearly with the number of available cores. Our software, HPC-REDItools, represents the first HPC resource specifically devoted to RNA-editing detection, allowing the analysis of individual RNA-seq samples in a few minutes. The software is available at https://github.com/BioinfoUNIBA/REDItools2.

## Implementation

RNA-editing events detected by REDItools are represented in a machine-readable tabular format as reported, for example, in Figure 1 in which each row is an editing site characterised by the chromosome name (column *Region*) and genomic coordinate (column *Position*). In addition, each editing position includes additional attributes, such as the reference nucleotide (column *Reference*), the list of observed substitutions (column *AllSubs*), the RNA editing frequency (column *Frequency*) (representing the editing level per site), the number and base count of reads supporting the position (columns *Coverage-q25* and *BaseCount[A,C,G,T]*, where q25 refers to the minimum quality score of a base), the strand (column *Strand* where 1 indicates strand plus, 0 strand minus and 2 strand not defined) and the mean quality of supporting nucleotides (column *MeanQ*).

**Figure 1.**
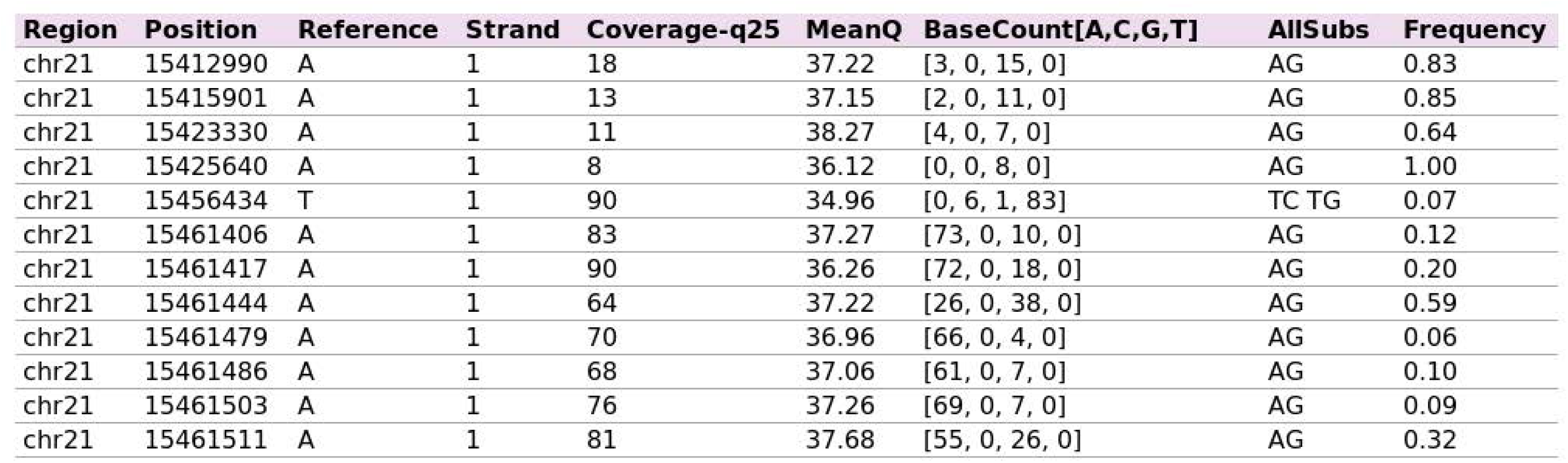
Example of the machine-readable tabular format used by REDItools to represent RNA-editing events.

The identification of RNA editing events in a classical RNA-seq experiment (including at least 50-60 million reads) through the first implementation of REDItools [13] can take up to 300 hours on a standard machine with a single core and 2GB of RAM. Consequently, REDItools are not suitable to analyse large datasets from public repositories (e.g. GTEx or TCGA). To overcome this issue, we have completely redesigned the main algorithm and adapted it to HPC infrastructures to benefit from multiple parallel cores.

Our novel software, called HPC-REDItools, introduce at least three big novelties over the previous implementation:

1. **Data loading optimization** the new code has been fully rewritten to optimise data loading and improve computational speed when launched in serial mode;
2. **Dynamic Interval Analysis** we have implemented a novel algorithm, called *Dynamic Interval Analysis* (DIA), to solve the problem of high-density genomic regions that improves the software in terms of workload balance;
3. **Parallelization** the current code is HPC-aware supporting the parallel analysis on thousands HPC nodes.

Below we describe in detail algorithmic novelties illustrating speed improvements over the original REDItools implementation.

### HPC-aware implementation

Original REDItools software is written in Python (version 2.7) and for its biological purpose (RNA editing detection) employs the *pysam* module, a wrapper of the widespread SAMtools package for reading and manipulating large raw BAM files containing multiple alignments of transcriptome reads onto a reference genome (composed by several sequences corresponding to chromosomes). REDItools can speed up the identification of editing events using multi-threading in which each thread (available core) analyses independently a complete chromosome or part of it. Such pseudo-parallelism is strongly limited because REDItools run on single machines and, thus, the number of useful processes and analyses is restricted to the number of available cores (typically in a range from 2 to 30 on modern CPUs). In addition, REDItools do not take into account the density of reads per chromosome and some processes may be more computationally stressed than others. HPC-REDItools, instead, have been designed to be highly-parallel in order to run on HPC infrastructures, as they are becoming more and more accessible to researchers worldwide, and take advantage of multiple computing nodes. HPC-REDItools are again written in Python (to increase portability and for continuity with the previous version) and makes use of mpi4py library [17] (version 2.0.0) that is the binding of the Message Passing Interface (MPI) standard library for the Python programming language. Such library enables the power of multi-node computing and gives access to point-to-point and collective communication primitives (e.g., send/receive, scatter/gather and son on) directly from native Python code. The general architecture of HPC-REDItools follows a simple master/slave template. As shown in Figure 2, the MPI program (yellow rectangle) takes in input the BAM file (see Fig. 2 (A)). A master process *M* splits the input whole genome into a set of genomic intervals *D* and, then, dispatches each interval to *n* free parallel slave processes by sending a *COMPUTE* message and until all processes are kept busy. Each slave process *S* (Fig. 2 (D)) performs the analysis on the assigned interval and produces an intermediate temporary file with candidate RNA-editing events (see Fig. 2 (E)). To promote process recycling, whenever a slave process *S* completes its analysis, it notifies the master process *M* by sending a *DONE* message which in turn assigns a new interval to *S* (if any). When all intervals have been analysed, the master process *M* sends a *FINISH* message to each slave process *S* notifying the end of the computation so that they can gracefully exit. A final procedure (Fig. 2 (F)) is implemented to collect all intermediate results (temporary files) and creates a single, unified output file with all potential RNA-editing events (Fig. 2 (G)). HPC-REDItools accept personalised intervals for an advanced and fine-grained control over the computation. However, we have implemented the DIA Algorithm (see Algorithm 1 in Section) which exploits the point-wise coverage of a given BAM file to return an optimal set of intervals and guarantee a more balanced workload across processes.

**Figure 2.**
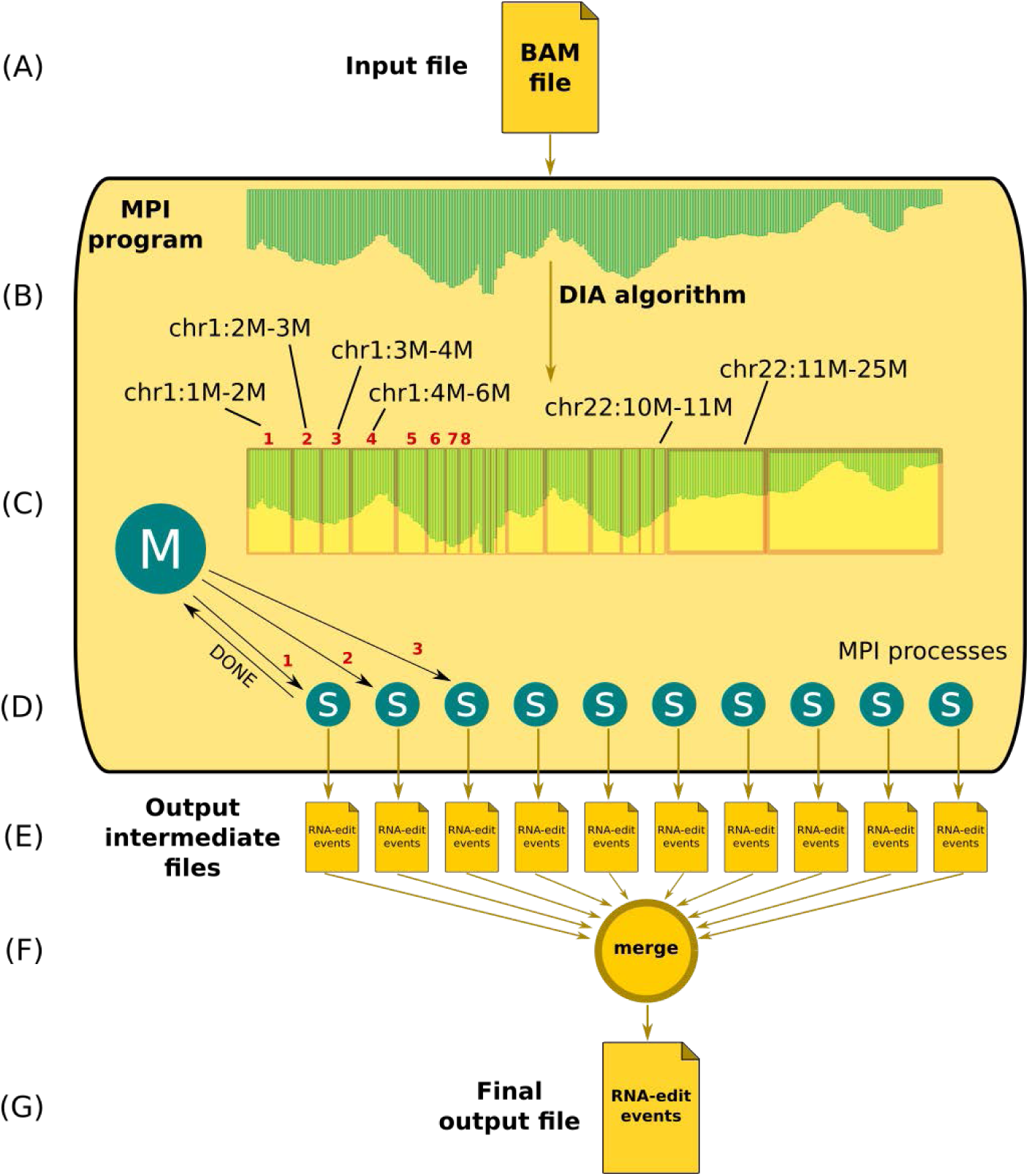
Layout of HPC-REDItools parallel implementation.

### Data loading optimization

A relevant novelty of HPC-REDItools consists in the optimization of the data loading which represented a bottleneck of the previous REDItools version. Such optimization improves the global speed per genomic interval and has a positive impact on parallelism. As explained above, the original REDItools implementation relies on the well-known Python module `Pysam` [18] (version 0.13) to extract and manipulate aligned reads in SAM/BAM format. To look at RNA editing events, REDItools inspect the entire genome position by position iteratively by invoking the `mpileup` function of Pysam to calculate statistics about transcriptome reads supporting each position. During the traversal, the `mpileup` function loads scanned reads as many times as their length, causing *l* disk accesses per read (where *l* is the read length) (see Figure 3, left). To overcome this limitation, individual genomic positions could be simply explored by traversing aligned reads sequentially with no need to use the `mpileup` function. Indeed, HPC-REDItools access the disk only once for each read: as a new genomic position is encountered, it dynamically loads from the disk all the novel mapping reads present in the input BAM file which start at the given position, caching the information as needed for future reuse (see Figure 3, right). Note that as soon as a read does not intersect the position currently under analysis, it is discarded and the corresponding memory freed, thus also contributing to a minimal memory footprint. Furthermore, the rate of disk accesses is decreased by a factor equal to *n* and the overall analysis time is dramatically reduced.

**Figure 3.**
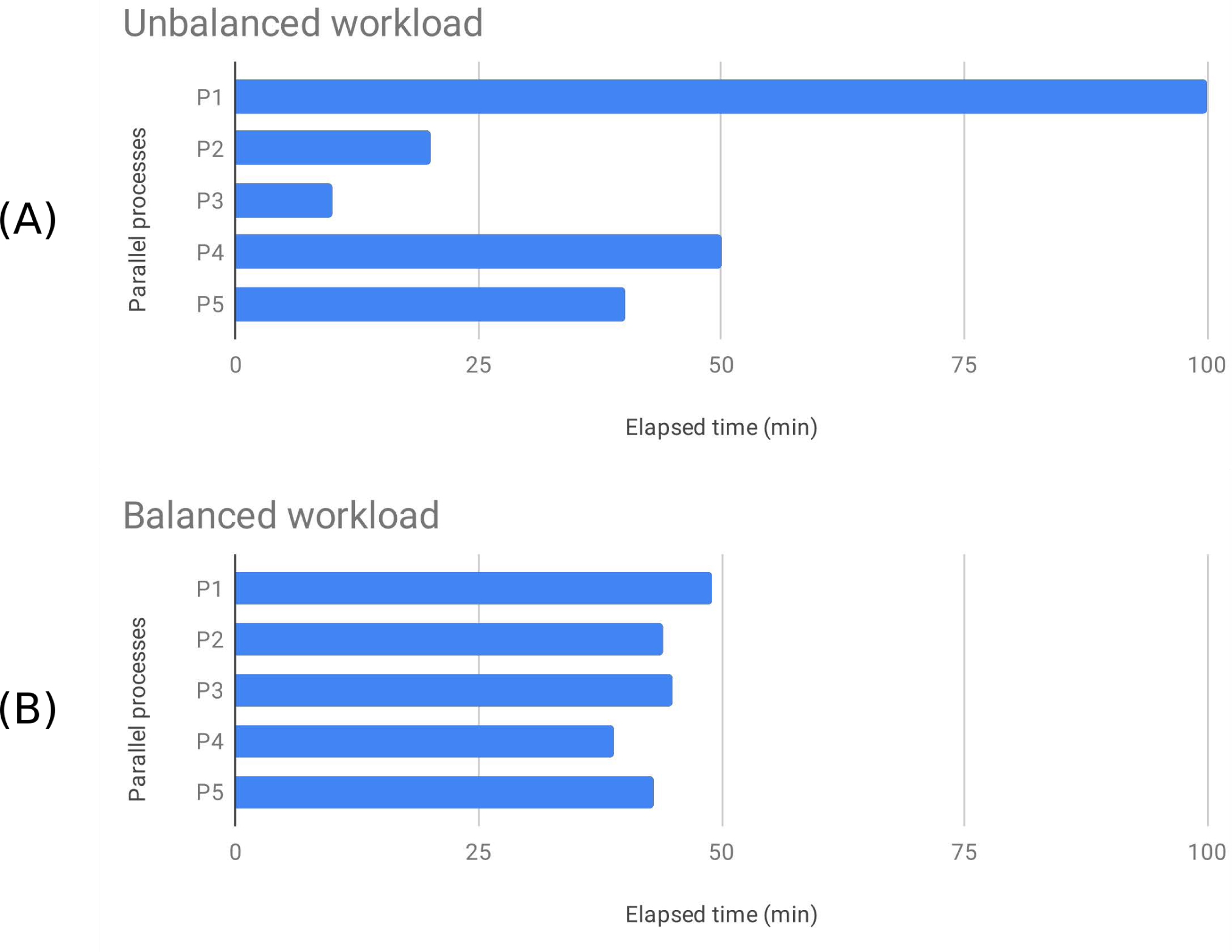
Data loading improvement. The figure shows the BAM traversal strategy in the original serial version of REDItools (left) and in HPC-REDItools (right). While in the previous version each read was loaded *n* times, in HPC-REDItools each read is loaded just once, thus reducing the overall bottleneck introduced by useless accesses to disk.

### Dynamic Interval Analysis

The parallelism of the original REDItools version consists in splitting the genome into *n* equal-sized intervals, where *n* is the number of processes specified at the program invocation. Genomic intervals of RNA-seq experiments are characterised by an extreme uneven coverage (density of mapped reads) and when treated equally result in an unbalanced workload across processes with sub-optimal performances (cfr. Fig. 4 (A) and (B)). In addition, splitting highly-variable coverage data into equal-sized intervals causes computational time bottlenecks with fast processes for low coverage intervals and very slow processes for high-density intervals. In the worst case, a unique process might monopolise the entire time slice allocated to the whole job because of an interval with an extremely high number of supporting reads. However, many biological data produced by high throughput sequencing technologies are characterised by a highly-variable coverage and, thus, ad-hoc solutions are needed.

**Figure 4.**
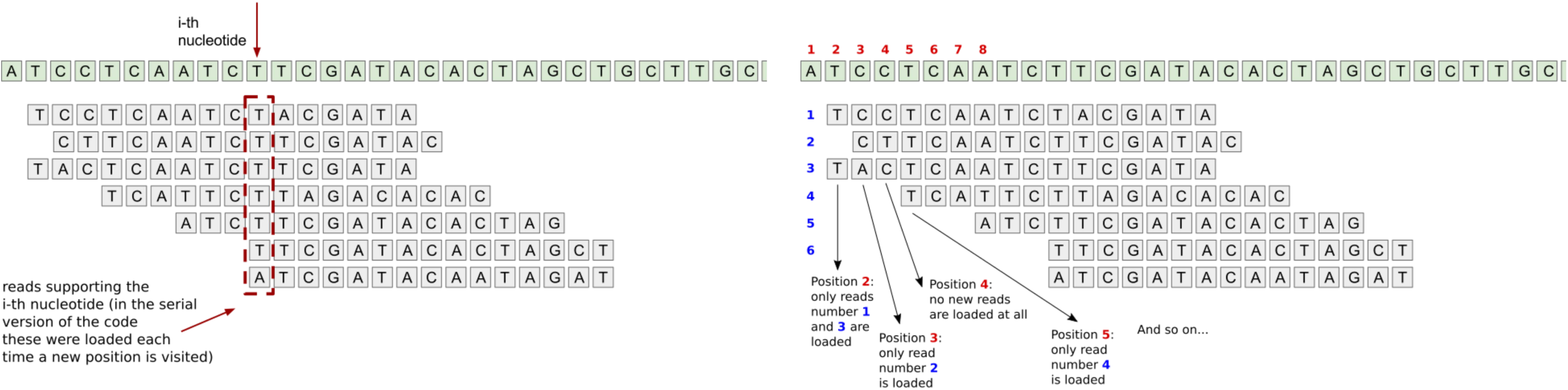
Unbalanced vs. balanced workload. Examples of the impact of different intelligent interval divisions on a system’s workload balance: when a naive interval division is employed, the system potentially suffers from unbalanced workload (A) which is instead overcome by our *Dynamic Interval Analysis* thanks to which the workload skyline becomes more homogeneous and all processes gain in fairness (B).

HPC-REDItools introduce a novel strategy called *Dynamic Interval Analysis* (DIA). Such strategy aims at finding an optimal interval division, which is able to guarantee a balanced processing time across processes while maintaining the number of intervals comparable to that of processes. Formally, the goal is to find a set *D* of *n* intervals (*n* fixed) such that the processing time *T* (*I*) of each interval *I* is approximately the same, that is:

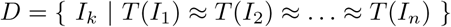

This is equivalent to say that *T* (*I*_*i*_) *≈ c*_*n*_, meaning that the time to process a given interval is constant (equal to *T* (*G*)*/n*, where *T* (*G*) is the total time required to analyse the whole genome *G* in a serial fashion). Since the function *T* is not known, it is necessary to find the best estimate 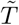 of *T* which is able to predict the execution time over an interval. First, we define the time needed to analyse an interval *I* as the sum of the single contributions over each position *i* in the interval, thus reducing the problem of estimating the processing time over single positions:

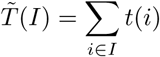

where *t*(*i*) is a function that estimates the processing time of the *i*-th position. For simplicity, we assume that the processing time to compute a certain position depends only on the number of reads supporting such position (i.e., known as *coverage*).

As shown in Algorithm 1, we first initialise the set *D* of dynamic intervals and then calculate the processing-time estimate *GC* for the whole genome (lines 3-6). We then calculate *AC* as the average coverage time of the ideal interval (line 7); this is the ideal constant processing time for an interval. Since intervals cannot overlap and are sets of contiguous positions, the algorithm starts at the first position of the genome and then proceeds by expanding the partial interval until a stop condition is reached. There are three possible stop conditions:

#### Algorithm 1 Dynamic Interval Analysis algorithm

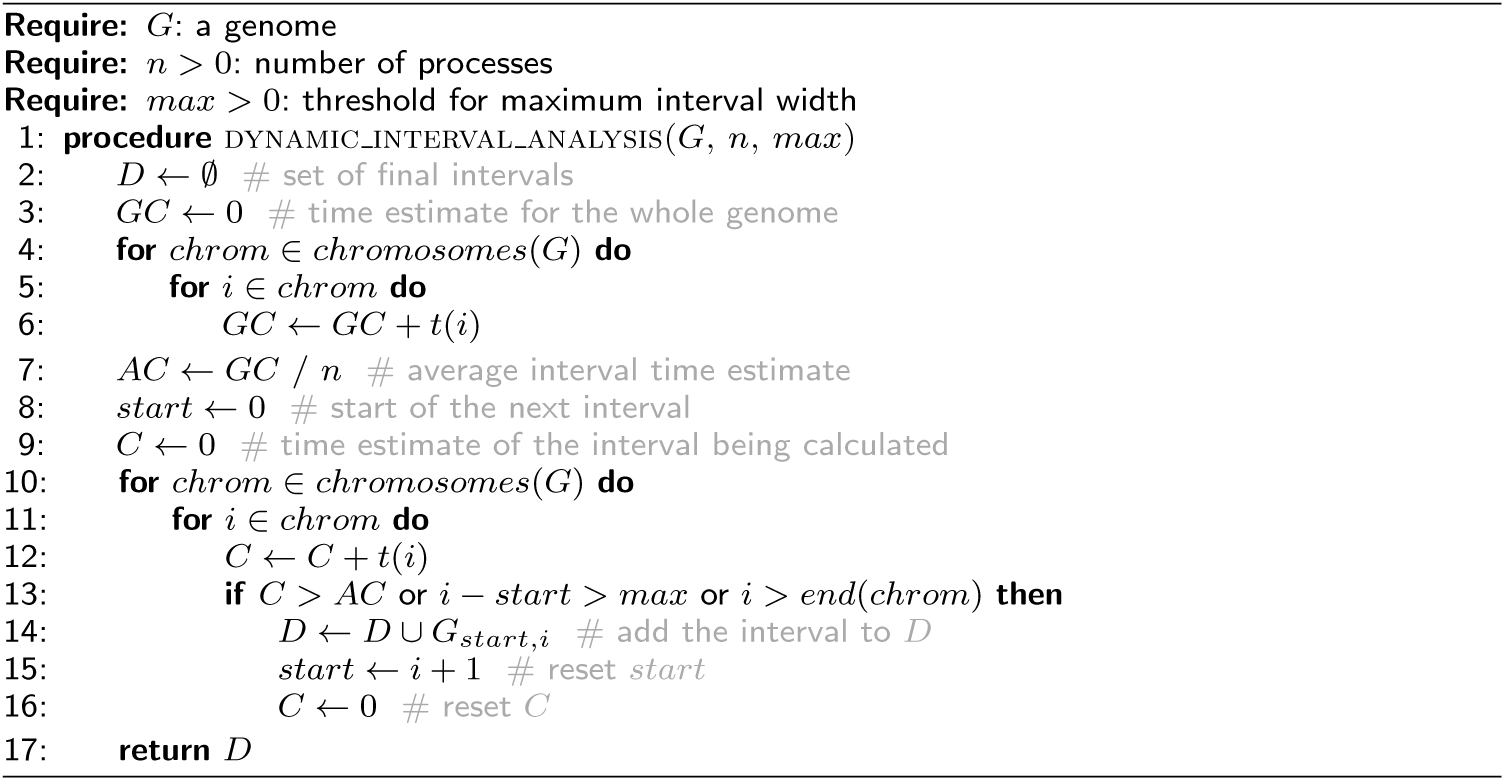

C1. **max-weight**: the weight of the interval under analysis exceeds the constant *AC* (to preserve workload balance);

C2. **max-width**: the width of the interval exceeds the desired threshold *max* (to encourage process recycling);

C3. **single-chromosome-span**: the end of a chromosome has been reached (since an interval cannot span across chromosomes).

### Finding time estimate

To find the best estimate for *t*, we selected a random sample of 1,000 equal-size intervals uniformly distributed across all human chromosomes, in order to take into account also short sequences (such as the mitochondrial genome, barely 16,571 base pairs in length). We analysed each interval with HPC-REDItools in serial mode (i.e., one single core) and calculated its average coverage. Figure 5 shows a log-log plot, where each (*x, y*) point represents an interval with average coverage *e*^*x*^ which has taken *e*^*y*^ seconds to be completed. As shown, the plot reveals two main clouds, the first representing intervals whose log average coverage is between 4 and 12 and the second which includes higher-coverage intervals. This plot is meaningful in the sense that it gives a pragmatic indication of the correlation between an interval coverage and the processing time it requires to analyse it. Since the analytical continuous function *t* is not known a priori, we tested several *t*, including constant, linear and polynomial functions. However, the plot suggests that the average time to elaborate a given interval correlates in a cubic manner with its mean coverage and the light-blue line in the figure represents the function which best fits the given discrete point distribution.

**Figure 5.**
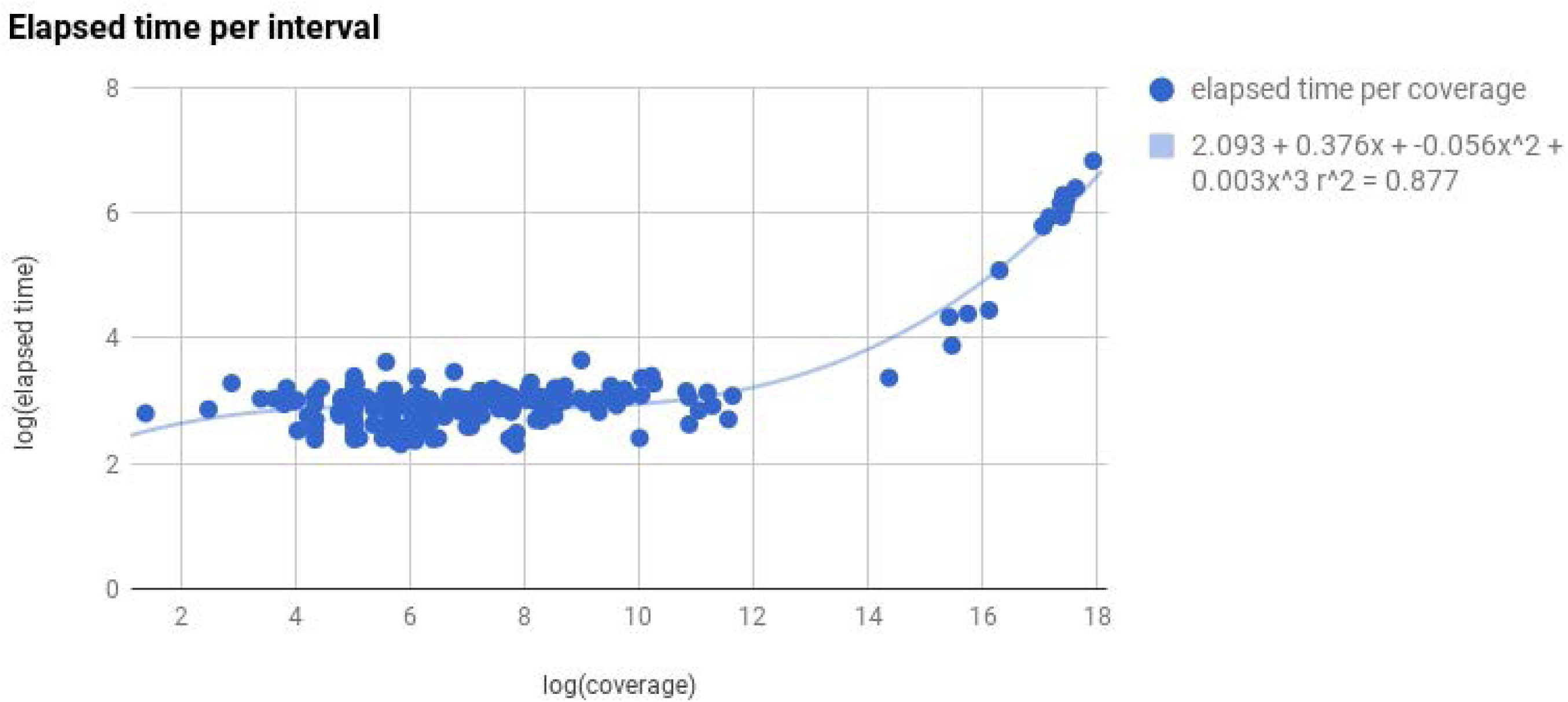
Correlation between coverage and processing time for a sample of 1,000 equal-size random intervals.

## Results

### Data loading optimization

To experimentally test the speed improvement between HPC-REDItools and REDI-tools, we created a dataset consisting of 10 RNA-seq samples randomly selected from the GTEx project. Both programs were launched on the same computer machine in order to predict RNA editing events occurring in the chromosome 21. In Figure 6 we report elapsed times for REDItools (in red) and HPC-REDItools (in blue). As shown, HPC-REDItools are on average 8 times faster than REDItools. This finding is quite interesting because enables the use of HPC-REDItools also to users with no access to HPC infrastructures and greatly speeds up the genome wide RNA editing detection.

**Figure 6.**
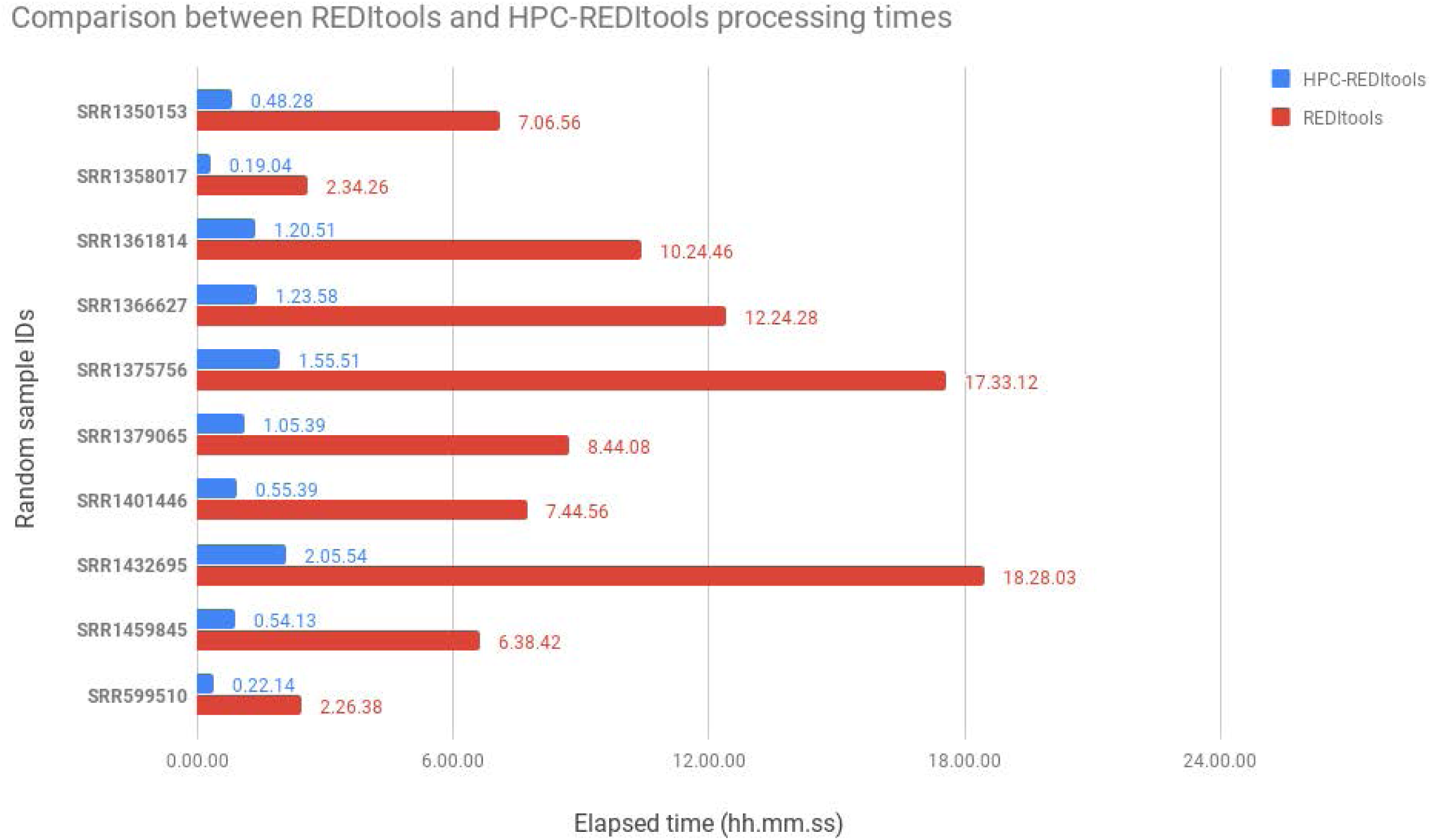
Comparison between REDItools and HPC-REDItools in terms of elapsed computing times when analysing 10 random samples downloaded from the GTEx repository. Comparison tests were performed on the same machine.

### Scaling analysis

To formally assess the scalability of HPC-REDItools on a High Performance Computing architecture, we generated a dataset comprising 200 **averaged-sized** samples from the GTEx repository and then we performed 7 experiments involving the analysis of 1, 2, 10, 20, 50, 100 and 200 samples, respectively. For each experiment we ran HPC-REDItools with a number of nodes equal to the number of selected samples (for example, for 10 samples we required 10 HPC nodes). All computations were executed on the Knight-Landing (KNL) partition of Marconi, a Tier-0 cluster available at CINECA, the Italian biggest non-profit organization offering support to national and international research projects as well as one the most powerful calculus infrastructures in Europe.^[3]^ Results are shown in Figure 7 (left). The x-axis reports the number of requested nodes while the y-axis indicates the achieved speed-up, defined as 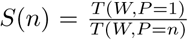 that is the fraction of the time required to perform some work *W* using 1 core over the time required to perform the same amount of work using *n* cores; for example,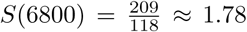. The optimal speed-up is *s* = 1, corresponding to the implementation that achieves the maximum parallelism possible. As shown in the figure, in which blue line corresponds to HPC-REDItools timings and red line to the optimal speed-up, HPC-REDItools achieves a very good scaling, demonstrating the ability to optimally exploit the computational power offered by a HPC infrastructure.

**Figure 7.**
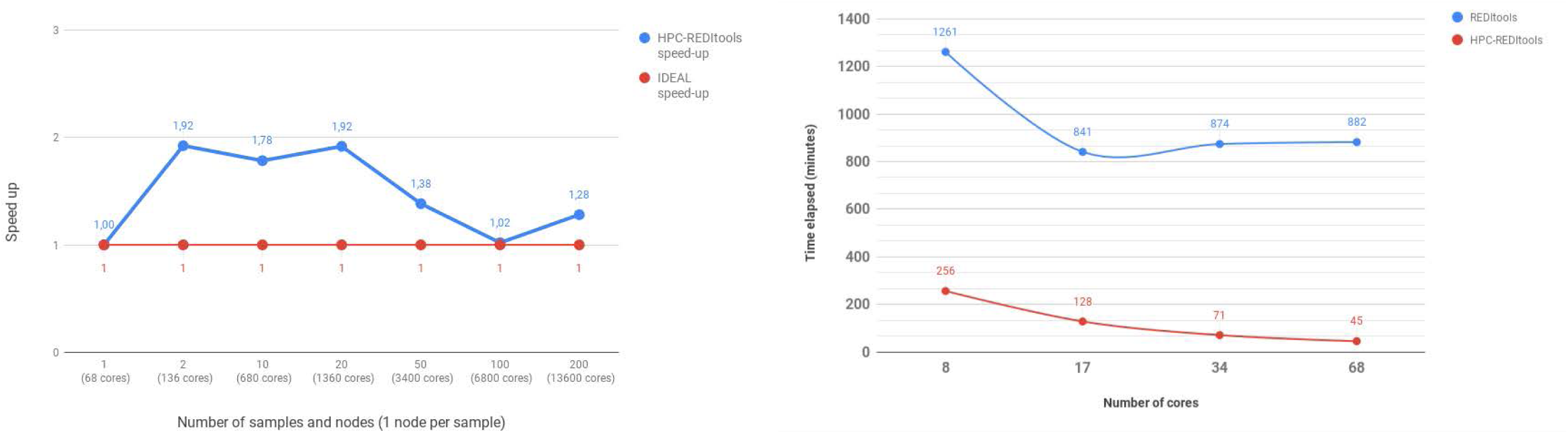
Scaling behaviour of HPC-REDItools: speedup (left) and comparison between the two versions of REDItools (right).

We then compared the scalability of REDItools and HPC-REDItools on an individual sample of 2.7 GB by varying the number of requested cores (*x* = 8, 17, 34, 68) on a single KNL node (see Figure 7, right).As a result, HPC-REDItools appeared about 8 times faster than the previous version enabling the RNA-editing detection in less than 1 hour. In contrast, REDItools reached a plateau at *x* = 17, due to the fact that it barely splits the whole genome into chromosome-wide regions, assigning each region to a different thread, thus wasting all the computational power coming from the additional cores.

## Conclusions

RNA editing is a co-/post-transcriptional phenomenon occurring in many organisms including animals and plants and has relevant biological implications. It can be detected employing RNA-seq data generated by high throughput sequencing technologies. However, as data volume increases, more powerful tools are required to analyse large number of samples in a time affordable way. In the present work we described HPC-REDItools, a HPC-aware tool for efficiently detect high-quality RNA-editing events from big data repositories on a HPC cluster. HPC-REDItools introduce at least three main algorithmic improvements over the previous version: i) high parallelism to employ the computational power available at High Performance Computing infrastructures; ii) optimised data loading that dramatically reduces computing time per genomic interval; iii) Dynamic Interval Analysis approach to improve workload balance across parallel processes. Our results indicate that HPC-REDItools are ready to analyse RNA editing in a variety of samples. Indeed, we plan to apply our software to explore the RNA editing landscape in large NGS datasets, thus providing a more reliable overview of the role of RNA editing in eukaryotic organisms.

### Availability and requirements

- Project name: HPC-REDItools
- Project home page: https://github.com/BioinfoUNIBA/REDItools2
- Operating system(s): Platform independent
- Programming language: Python *≥*2.7
- Other requirements: None
- License: GPL-3.0

## Acknowledgements

We would like to thank Claudio Lo Giudice and other colleagues from the RNA editing community for helpful suggestions.

## Author’s contributions

TF has re-implemented REDItools and is the principal writer of the manuscript; SG has tested the software and contributed to the writing of the manuscript; NS has contributed to the testing of the parallel version of HPC-REDItools; IT, MAD and GP have equally contributed to the writing of the manuscript; GC, EP and TC have guided the research and contributed to the writing of the manuscript. All authors read and approved the final manuscript.

## Availability of data and materials

Source code and examples are available on GitHub.

## Ethics approval and consent to participate

Not applicable.

## Consent for publication

Not Applicable.

## Footnotes

[1] https://cancergenome.nih.gov/

[2] http://gtexportal.org/

[3] Nodes in the KNL partition of Marconi mount 68-core Intel(R) Knights Landing @ 1.40GHz processors.

